# Phylogenomic analysis of the understudied *Neisseriaceae* species reveals a poly- and paraphyletic *Kingella* genus

**DOI:** 10.1101/2022.10.13.512104

**Authors:** Daniel P. Morreale, Joseph W. St Geme, Paul J. Planet

## Abstract

Taxonomic classification and phylogenetic analysis of the *Neisseriaceae* family has focused on the pathogens *Neisseria meningitidis* and *Neisseria gonorrhoeae*. Less is known about the relationships of commensal *Neisseria* species and other *Neisseriaceae* genera, raising the possibility that the phylogeny of this family may not agree with taxonomy. In this study we used available nucleotide sequences and a phylogenetic approach to assess the *Kingella* genus and its relatives. We found that this genus is both paraphyletic and polyphyletic. *Kingella potus* is more closely related to *Neisseria bacilliformis* than other *Kingella* species. The *Alysiella* and *Simonsiella* genera form a distinct clade within the *Kingella* genus that is closely related to the pathogens *K. kingae* and *K. negevensis*. We find a phylogenetic relationship between *<u>C</u>onchiformibius, <u>A</u>lysiella, <u>S</u>imonsiella*, and *<u>K</u>ingella*, which we name the CASK clade. Finally, we define the gene sets that differentiate each genus of the CASK clade from one another and from the rest of the *Neisseriaceae* family.

**Importance:** Understanding the evolutionary relationships between the species in the *Neisseriaceae* has been a persistent challenge in bacterial systematics due to high recombination rates in these species. Previous studies of this family have focused on *N. meningitidis* and *N. gonorrhoeae*. However, previously understudied *Neisseriaceae* species are gaining new attention, with *K. kingae* now recognized as a common human pathogen and with *Alysiella* and *Simonsiella* being unique in the bacterial world as multicellular organisms. A better understanding of the genomic evolution of the *Neisseriaceae* can lead to identification of the specific genes and traits that characterize the remarkable diversity of this family.

## Introduction

Defining taxonomic relationships is a cornerstone of microbial systematics, but in species that reproduce asexually and trade genetic information through active horizontal gene transfer across the species boundary, defining the relationships between species can be challenging. As in many taxonomic disciplines, the relationships amongst and within species have traditionally been established using bacterial cell morphology and other phenotypic properties. With the advent of molecular techniques such as DNA-DNA hybridization, 16S rRNA gene sequencing, multilocus sequence typing (MLST), and matrix-assisted laser desorption/ionization time of flight (MALDI-TOF), this system gave way to the so called polyphasic classification strategy that seeks to consider both phenotypic and molecular data (1, 2). As whole genome sequences became readily available, a newer version of this system, taxonomogenomics, seeks to incorporate this rich genomic data into taxonomical descriptions (3–6). This approach also has the benefit of being explicitly phylogenetic, providing a non-arbitrary approach to classification (5, 7, 8). While there is still controversy about how best to integrate phenotypic and phylogenomic data in bacterial taxonomy, there is a general consensus that taxonomy should be reflective of the evolutionary relationships and that monophyly is a desirable characteristic of taxonomic groups.

The *Neisseriaceae* family is a group of Gram-negative, β-proteobacteria that currently includes 20 genera: *Neisseria, Wielerella, Vitreoscilla, Uruburuella, Stenoxybacter, Snodgrassela, Simonsiella, Rivicola, Prolinoborus, Paralysiella, Morococcus, Kingella, Eikenella, Crenobacter, Craterilacuibacter, Conchiformibius, Bergeriella, Aquella, Amantichitinum*, and *Alysiella* (9–11). The majority of prior work devoted to understanding the taxonomic relationships between these genera has focused primarily on the pathogenic *Neisseria* species, *N. meningitidis* and *N. gonorrhoeae* (2, 8, 12–17). Early work using 16S rRNA failed to properly resolve species boundaries, resulting in several polyphyletic genera (18). In addition, pervasive horizontal gene transfer (HGT) and recombination in the *Neisseriaceae* further increases taxonomic ambiguity, with clonal lineages punctuated by large recombination events between distant relatives (14, 19). HGT results in the formation of ill-defined species with ambiguous species boundaries (20). To help resolve this ambiguity, recent studies of the commensal *Neisseria* species employed core genome MLST (cgMLST) and proposed genus specific cut-offs for defining species using genome relatedness indices (8, 19, 20). However, these studies did not comprehensively address other polyphyletic genera within the *Neisseriaceae*.

The *Kingella* genus includes five taxa: *K. potus, K. oralis, K. denitrificans, K. negevensis*, and *K. kingae* (21–27). Recently, two novel species have been proposed, namely *K. bonacorsii and K. pumchi* (28, 29). All of the *Kingella* species are typically associated with the oral or oropharyngeal microbiome, a common niche for other *Neisseriaceae*. Isolates of *K. kingae, K. negevensis*, and *K. potus* have all been associated with invasive disease in immunocompetent individuals (23, 25, 30). In contrast, *K. oralis* and *K. denitrificans* are present in dental plaque and in gingival and periodontal samples and only rarely cause invasive disease, typically in immunocompromised individuals (26, 31, 32). *Kingella* species are oxidase positive, catalase negative, rods and typically form short chains (33).

A review of published phylogenetic relationships among the commensal *Neisseria* species reveals a high level of variability in the proposed taxonomic relationships, primarily owing to the genes selected and the method used to perform the analysis (15, 34–36). In many of these studies the *Kingella* genus is polyphyletic, and the genera that interrupt a monophyletic clade with *Kingella* species differ in different published analyses (for examples see 24, 33, 34, 37, 38). Most isolates used for defining taxonomic nomenclature in this genus were collected prior to pervasive whole genome sequencing and were assigned to *Kingella* as a result of phenotypic and/or minimal sequence-based analyses. Additionally, as the frequency of culture-dependent and culture-independent microbiome studies have become more prevalent, the number and diversity of formally accepted species in the *Neisseriaceae* have increased (13). Here, we seek to reassess the relatedness of the five species in the *Kingella* genus using whole genomes and a phylogenomic approach.

## Results

### *Kingella potus* is distinct from the other species in the *Kingella* genus

To examine the *Kingella* species, 16S rRNA gene sequences for isolates from the *Neisseriaceae* were collected for phylogenetic reconstruction (Fig. 1). By 16S rRNA relatedness alone, it is clear that a major driver of the polyphyletic structure of *Kingella* is *K. potus*. This species was recovered from an infected wound caused by an animal bite and is more closely related to *Neisseria bacilliformis* than it is to any currently described *Kingella* species (25). Phenotypically, *K. potus* and *N. bacilliformis* are characterized for a limited number of characteristics, though both species are non-hemolytic, indole-negative coccobacilli that produce acid from glucose, maltose, and fructose (Table S3).

**Figure 1.**
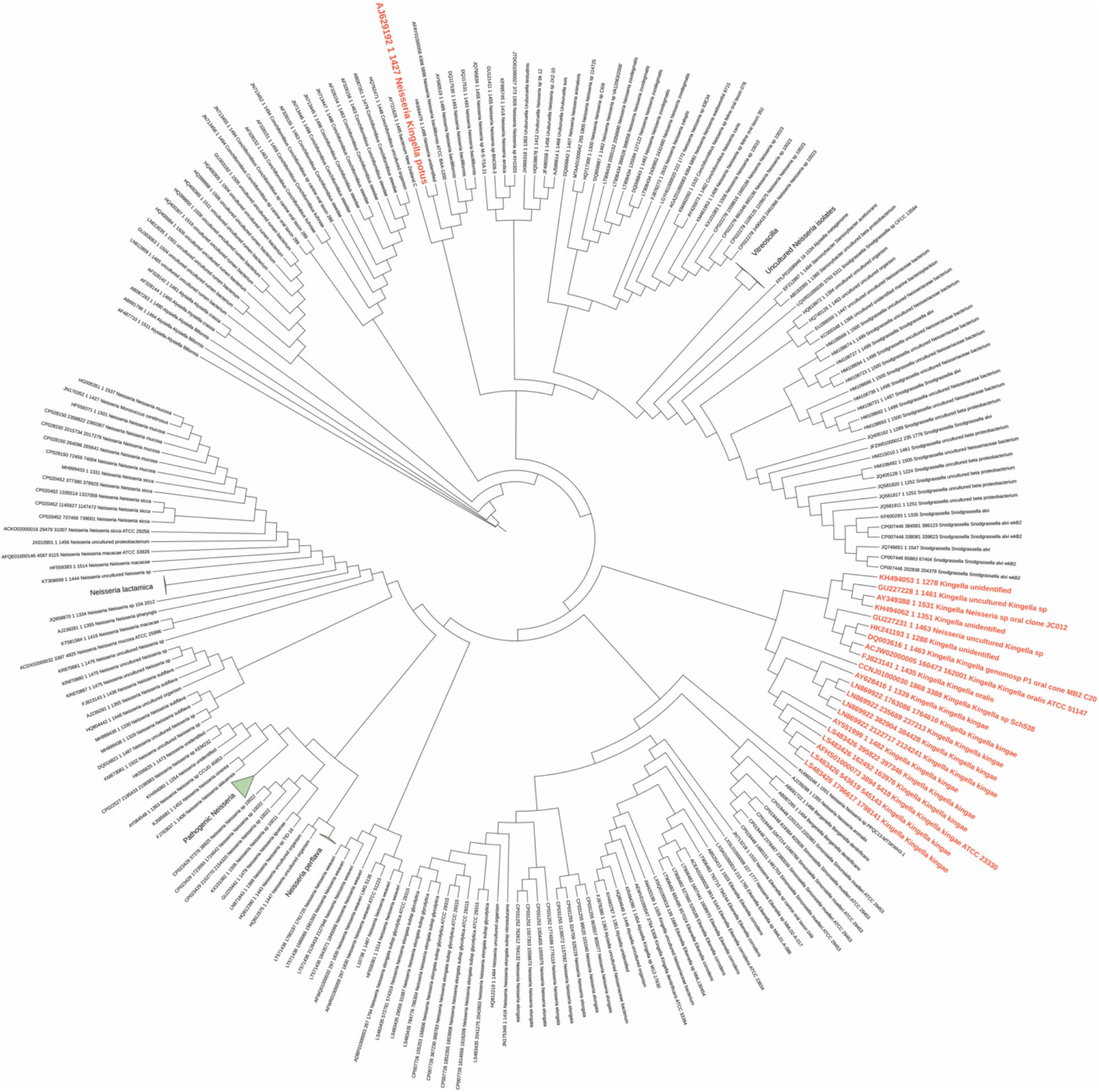
High quality 16S rRNA sequences were downloaded from the SILVA database for the family Neisseriaceae (taxid: 408). Sequences were aligned with MAFFT, and a maximum likelihood phylogeny was constructed using RAxML. Species that belong a species in the genus *Kingella* are colored red. By 16S rRNA relatedness, *Kingella* forms a polyphyletic clade with *K. potus* falling outside of the rest of the genus. *K. potus* forms a clade with the oral-associated *Neisseria bacilliformis*. Bootstrap values greater than 70% are shown.

To better understand the relationship between *K. potus* and *N. bacilliformis*, we calculated average nucleotide identity (ANI) and digital DNA-DNA hybridization (dDDH) between the type strains of these species. The ANI between *K. potus* and *N. bacilliformis* was calculated at 86.75%, below the established species cut-off of 95% (39). However, the ANI between *K. potus* and *N. bacilliformis* is appreciably greater than the ANI calculated between *K. potus* and other *Kingella* species, which ranges between <75% (*K. kingae*) and 78.8% (*K. denitrificans*) (Table 1, Fig. 2). The calculated dDDH between these species falls below species thresholds at 30% (95% confidence interval 27.6-32.5%) (Table 2).

**Table 1.** Average ANI scores calculated among taxa of interest. ANI scores were calculated using FastANI and are shown only if the ANI >75%. Empty cells denote data excluded from the table.

|  | *N.*<br>*bacilliformis* | *K.*<br>*denitrificans* | *K.*<br>*oralis* | *K.*<br>*negevensis* | *K.*<br>*kingae* | *N.*<br>*elongata* | *A.*<br>*crassa* | *A.*<br>*filiformis* | *S.*<br>*muelleri* |
| --- | --- | --- | --- | --- | --- | --- | --- | --- | --- |
| *K. potus* | 86.75 | 78.88 | 78.74 | <75 | <75 | 80.7 | - | - | - |
| *K. bonacorsii* | - | 79.84 | 96.41 | 78.06 | 77.9 | 78.76 | 77.98 | 77.87 | 77.14 |
| *A. crassa* | - | 77.81 | 77.96 | 79.37 | 79.74 | <75 | 100 | 85.43 | 79.79 |
| *A. filiformis* | - | 77.81 | 77.96 | 79.37 | 79.74 | <75 | 85.43 | 100 | 78.75 |
| *S. muelleri* | - | <75 | 76.9 | 79.16 | 79.96 | <75 | 79.79 | 78.75 | 100 |

**Figure 2.**
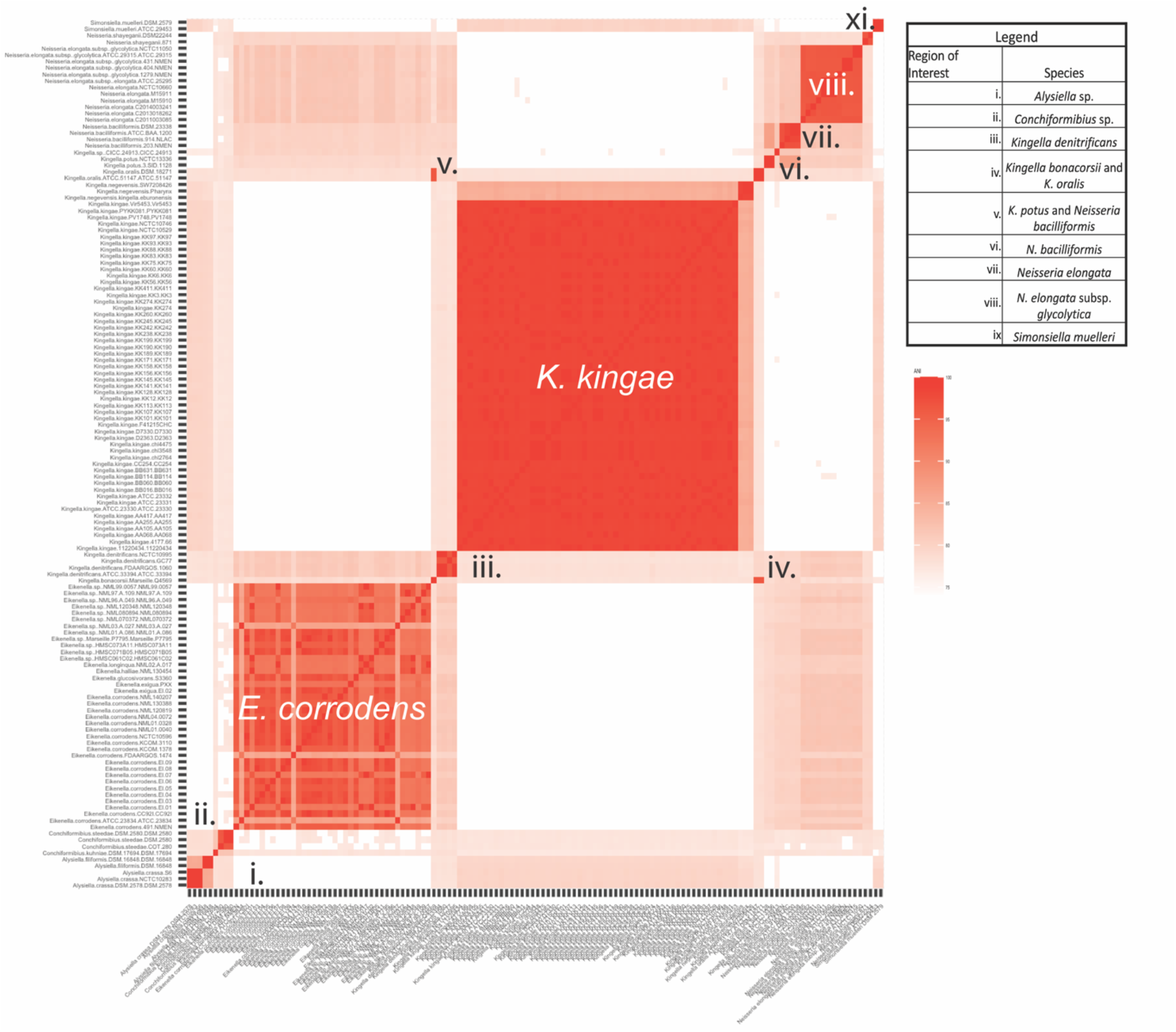
ANI scores comparing sequenced isolates from species in the *Neisseriaceae* most closely related to the genus *Kingella*. ANI scores were calculated between publicly available genomes downloaded from NCBI using FastANI. Several regions of interest are indicated on the heatmap, including the species delimiters, and regions of interest comparing *K. bonacorsii* and *K. oralis* (iv) and *K. potus* and *N. bacilliformis* (v). The resulting ANI matrix was rendered as a heatmap using R.

**Table 2.** dDDH values of *K. potus* calculated against closely related taxa. dDDH was calculated using TYGS using each of the possible algorithms, as well as the difference in G+C content between *K. potus* and each of the subject type strains.

| Query strain | Subject strain | dDDH<br>(d0, in %) | C.I.<br>(d0, in %) | dDDH<br>(d4<br>, in %) | C.I.<br>(d4, in %) | dDDH<br>(d6, in %) | C.I. (d6, in %) | G+C content<br>difference (in %) |
| --- | --- | --- | --- | --- | --- | --- | --- | --- |
| *K. potus*<br>NCTC13336 | *N. bacilliformis*<br>ATCC BAA-1200 | 45.6 | [42.2 -<br>49.0] | 30 | [27.6 -<br>32.5] | 41 | [38.0 -<br>44.0] | 1.86 |
| *K. potus*<br>NCTC13336 | *N. elongata* subsp.<br>*glycolytica* ATCC 29315 | 20.3 | [17.1 -<br>23.9] | 22.5 | [20.2 -<br>25.0] | 19.7 | [17.0 -<br>22.8] | 3.71 |
| *K. potus*<br>NCTC13336 | *K. negevensis*<br>Sch538 | 12.9 | [10.2 -<br>16.2] | 23.5 | [21.2 -<br>25.9] | 13.3 | [10.9 -<br>16.0] | 12.24 |

### The *Kingella* genus is paraphyletic

To further define the relationships among *Kingella* and closely related species, whole genome sequences for select species in the *Neisseriaceae* were downloaded from NCBI. As 16S rRNA gene sequences have been historically used to define species in this group, all following analyses were limited to species most closely related to *Kingella* by 16S rRNA gene phylogeny (Fig. 1). This group includes *Neisseria elongata, Neisseria sheyganii, N. bacilliformis, Eikenella corrodens, Alysiella filiformis, Alysiella crassa, Simonsiella muelleri, Conchiformibius steedae, K. oralis, K. bonacorsii, K. potus, K. denitrificans, K. kingae*, and *K. negevensis. N. elongata*, *N. sheyganii*, and *E. corrodens* were included as outgroups for these analyses. In total, 134 whole genome sequences were downloaded from NCBI and annotated with Prokka.

To minimize the impact of HGT and variability in the pangenome, Roary was used to identify the core genome (40). Core genes were clustered at the 75%, 80%, 85%, 90%, and 95% sequence similarity level and were limited to only those clusters of orthologous groups (COGs) found in all genomes. Core gene alignments were generated and used to reconstruct maximum-likelihood phylogenies (Fig. 3). Both the Robinson-Foulds distance and normalized Nye Similarity metrics were used to quantify tree similarity. Each phylogeny was highly congruent with the others based on these metrics, and each included a polyphyletic clade for the genus *Kingella* (Fig. 3B and C). We therefore proceeded with the lowest similarity cutoff of 75% for further analyses (Fig. 3A). In each of these analyses, *<u>C</u>onchiformibius* spp., *<u>A</u>lysiella* spp., *<u>S</u>imonsiella* spp, and *<u>K</u>ingella* spp. (with the exception of *K. potus*) form a closely related group that we will refer to as the CASK clade. Phenotypic data on the species in the CASK clade are extremely limited, limiting the utility of phenotypic properties in understanding relationships within the group (Table S3).

**Figure 3.**
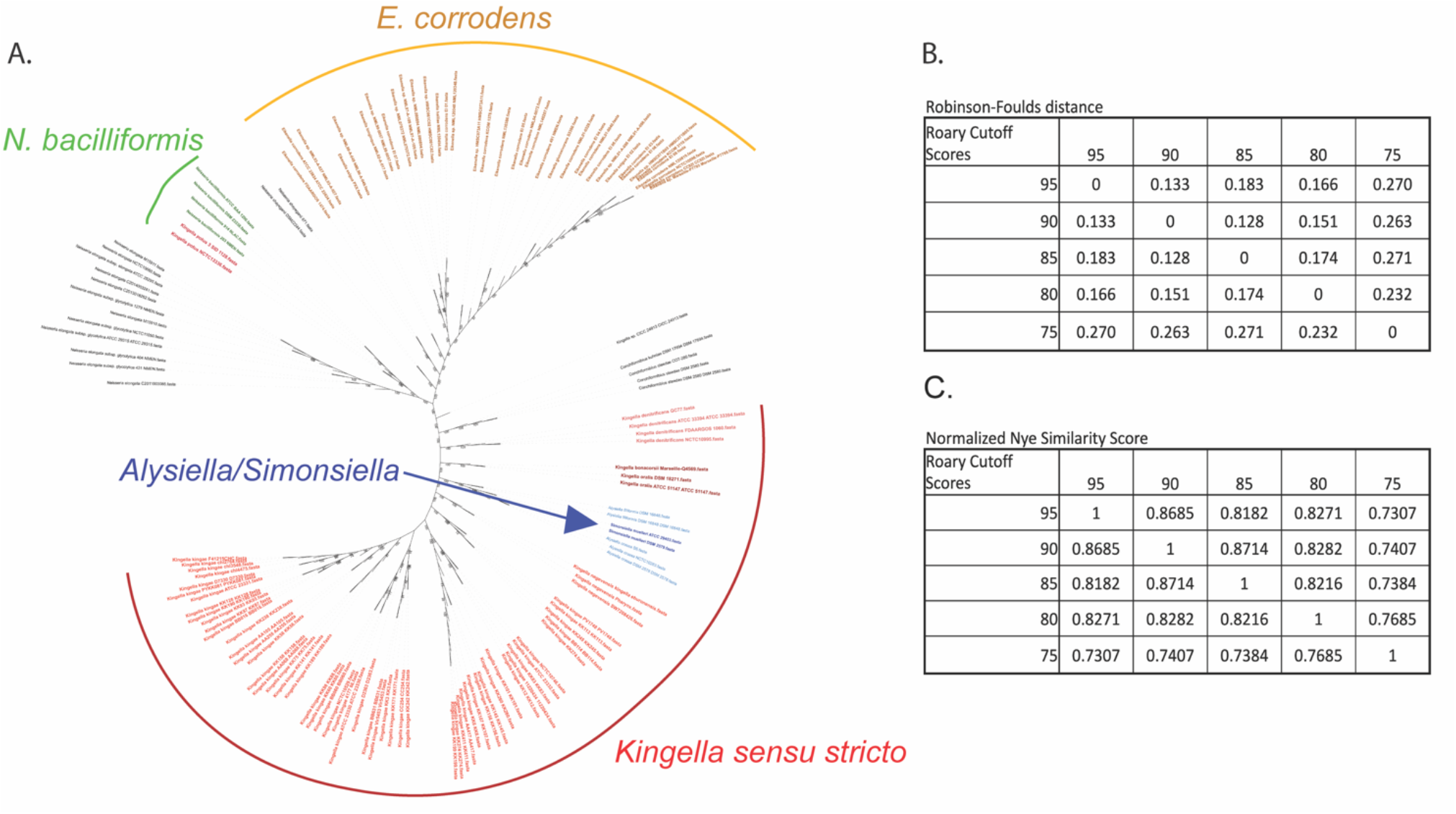
**A**. The core genome of the strains from the 13 species most closely related to *Kingella kingae* was determined using Roary with a minimum homology cutoff of 75%. The core genomes, which encompass approximately two million sites, were aligned with MAFFT and used to reconstruct a maximum likelihood phylogeny. As above, *K. potus* is only distantly related to the rest of the species in the genus *Kingella*. Additionally, the genera *Alysiella* and *Simonsiella* fall within the *Kingella* clade, suggesting the core genome of the species in each of these genera is very closely related to that of *Kingella*. Bootstraps greater than 70% are shown. **B-C**. The Robinson-Foulds and Normalized Nye similarity score congruency metrics were used to quantify changes in tree architecture between phylogenies reconstructed using core genome alignments. The Nye similarity score was normalized against the number of possible rearrangements given the number of leaves in the tree.

Based on a whole genome approach, *Alysiella* and *Simonsiella* form a clade within the *Kingella* genus (Fig. 3A), dividing *K. oralis* and *K. denitrificans* from *K. negevensis* and *K. kingae*. ANI, dDDH, and G+C% were calculated for type strains of *Kingella, Alysiella*, and *Simonsiella* and are shown in Tables 1 and 3. All of these overall genome relatedness indices (OGRI) support a close relationship between the species of these three genera.

**Table 3.**
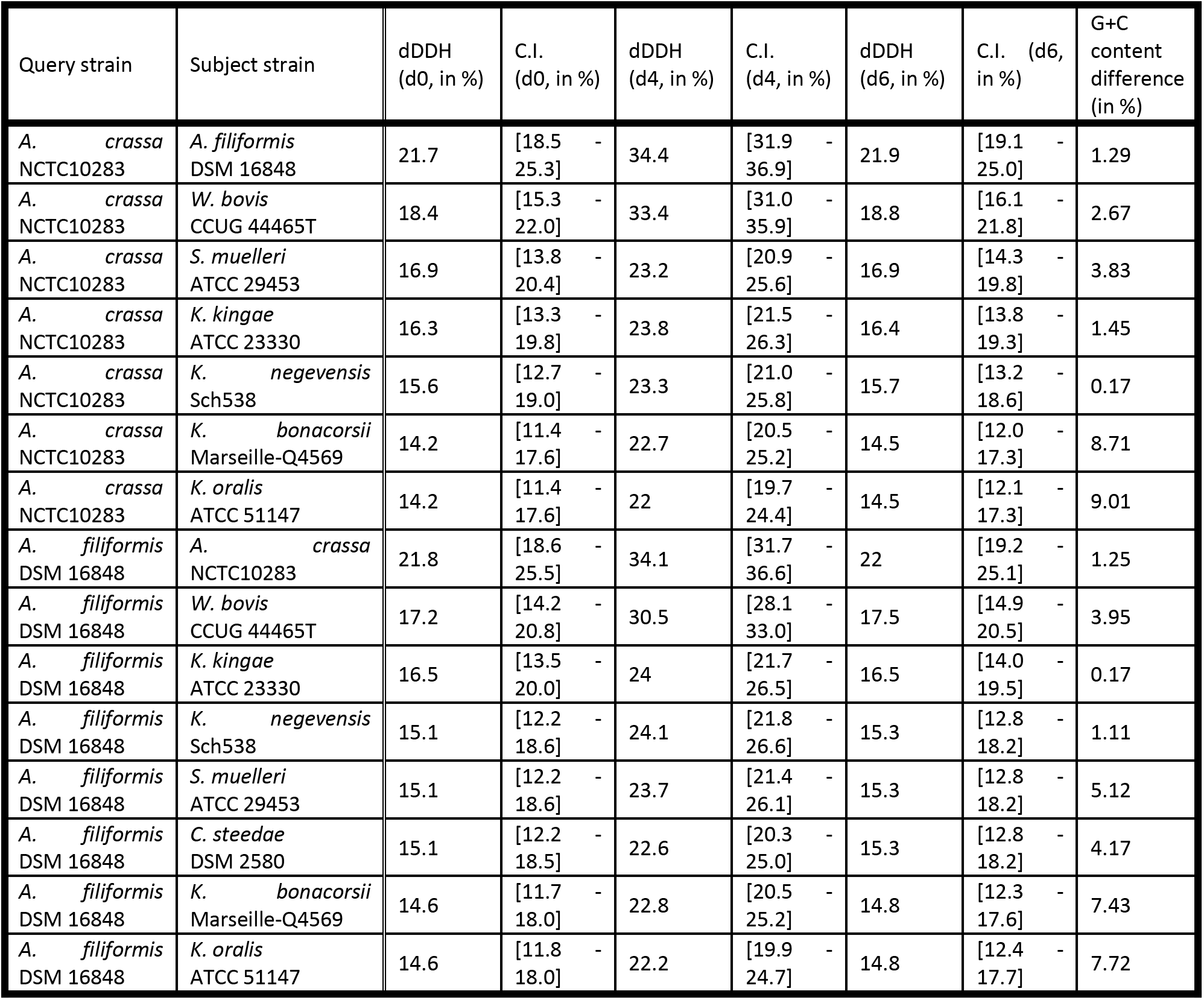
OGRI of *Kingella, Alysiella*, and *Simonsiella*. dDDH was calculated using TYGS using each of the possible algorithms, as well as the difference in G+C content between type strains of *Alysiella* or *Simonsiella* species and each of the closest related subject type strains.

| Query strain | Subject strain | dDDH<br>(d0, in %) | C.I.<br>(d0, in %) | dDDH<br>(d4, in %) | C.I.<br>(d4, in %) | dDDH<br>(d6, in %) | C.I. (d6,<br>in %) | G+C<br>content<br>difference<br>(in %) |
| --- | --- | --- | --- | --- | --- | --- | --- | --- |
| *A. crassa*<br>NCTC10283 | *A. filiformis*<br>DSM 16848 | 21.7 | [18.5 -<br>25.3] | 34.4 | [31.9 -<br>36.9] | 21.9 | [19.1 -<br>25.0] | 1.29 |
| *A. crassa*<br>NCTC10283 | *W. bovis*<br>CCUG 44465T | 18.4 | [15.3 -<br>22.0] | 33.4 | [31.0 -<br>35.9] | 18.8 | [16.1 -<br>21.8] | 2.67 |
| *A. crassa*<br>NCTC10283 | *S. muelleri*<br>ATCC 29453 | 16.9 | [13.8 -<br>20.4] | 23.2 | [20.9 -<br>25.6] | 16.9 | [14.3 -<br>19.8] | 3.83 |
| *A. crassa*<br>NCTC10283 | *K. kingae*<br>ATCC 23330 | 16.3 | [13.3 -<br>19.8] | 23.8 | [21.5 -<br>26.3] | 16.4 | [13.8 -<br>19.3] | 1.45 |
| *A. crassa*<br>NCTC10283 | *K. negevensis*<br>Sch538 | 15.6 | [12.7 -<br>19.0] | 23.3 | [21.0 -<br>25.8] | 15.7 | [13.2 -<br>18.6] | 0.17 |
| *A. crassa*<br>NCTC10283 | *K. bonacorsii*<br>Marseille-Q4569 | 14.2 | [11.4 -<br>17.6] | 22.7 | [20.5 -<br>25.2] | 14.5 | [12.0 -<br>17.3] | 8.71 |
| *A. crassa*<br>NCTC10283 | *K. oralis*<br>ATCC 51147 | 14.2 | [11.4 -<br>17.6] | 22 | [19.7 -<br>24.4] | 14.5 | [12.1 -<br>17.3] | 9.01 |
| *A. filiformis*<br>DSM 16848 | *A. crassa*<br>NCTC10283 | 21.8 | [18.6 -<br>25.5] | 34.1 | [31.7 -<br>36.6] | 22 | [19.2 -<br>25.1] | 1.25 |
| *A. filiformis*<br>DSM 16848 | *W. bovis*<br>CCUG 44465T | 17.2 | [14.2 -<br>20.8] | 30.5 | [28.1 -<br>33.0] | 17.5 | [14.9 -<br>20.5] | 3.95 |
| *A. filiformis*<br>DSM 16848 | *K. kingae*<br>ATCC 23330 | 16.5 | [13.5 -<br>20.0] | 24 | [21.7 -<br>26.5] | 16.5 | [14.0 -<br>19.5] | 0.17 |
| *A. filiformis*<br>DSM 16848 | *K. negevensis*<br>Sch538 | 15.1 | [12.2 -<br>18.6] | 24.1 | [21.8 -<br>26.6] | 15.3 | [12.8 -<br>18.2] | 1.11 |
| *A. filiformis*<br>DSM 16848 | *S. muelleri*<br>ATCC 29453 | 15.1 | [12.2 -<br>18.6] | 23.7 | [21.4 -<br>26.1] | 15.3 | [12.8 -<br>18.2] | 5.12 |
| *A. filiformis*<br>DSM 16848 | *C. steedae*<br>DSM 2580 | 15.1 | [12.2 -<br>18.5] | 22.6 | [20.3 -<br>25.0] | 15.3 | [12.8 -<br>18.2] | 4.17 |
| *A. filiformis*<br>DSM 16848 | *K. bonacorsii*<br>Marseille-Q4569 | 14.6 | [11.7 -<br>18.0] | 22.8 | [20.5 -<br>25.2] | 14.8 | [12.3 -<br>17.6] | 7.43 |
| *A. filiformis*<br>DSM 16848 | *K. oralis*<br>ATCC 51147 | 14.6 | [11.8 -<br>18.0] | 22.2 | [19.9 -<br>24.7] | 14.8 | [12.4 -<br>17.7] | 7.72 |

Additionally, we calculated pairwise ANI scores for each of the genomes listed above. The ANI scores confirm the majority of the proposed species classifications except for *K. bonacorsii*, an unclassified *Kingella* isolate that was recently proposed to be a new species (28). This isolate was recovered from a gingival sample, and the authors report a 16S rRNA similarity of 98.7%, ANI of 95.83%, and dDDH of 63.6% relative to *K. oralis* UB-38, as well as non-identification by MALDI-TOF mass-spectrometry (28). We replicated each of the genomic calculations reported as well as additional dDDH calculations using alternative formulae and found that choice of formula is essential, as only *d4* yielded a dDDH below the accepted values for species cutoffs (Tables 1 and 4, Fig. 2) (39, 41).

**Table 4.** COGs responsible for the unification of *Kingella s.s*. in the accessory genome.

| Last Common Ancestor *Alysiella*, *Simonsiella*, and *Kingella s.s.* | COG | State | p_0<sup>1</sup> | p_1<sup>2</sup> | Last Common Ancestor of *Kingella s.s.* | State | p_0<sup>1</sup> | p_1<sup>2</sup> | COG name<sup>3</sup> | Predicted Function<sup>3</sup> |
| --- | --- | --- | --- | --- | --- | --- | --- | --- | --- | --- |
|  | 742 | 1 | 0.15414 | 0.84586 |  | 0 | 0.98197 | 0.01803 | group_2532 | Protein IscX |
|  | 766 | 1 | 0.02993 | 0.97007 |  | 0 | 0.97924 | 0.02076 | group_2994 | hypothetical protein |
|  | 771 | 1 | 0.02993 | 0.97007 |  | 0 | 0.97924 | 0.02076 | *purU* | hypothetical protein |
|  | 923 | 1 | 0.24673 | 0.75327 |  | 0 | 0.97821 | 0.02179 | group_3650 | Sulfur carrier protein TusA |
|  | 3159 | 1 | 0.06887 | 0.93113 |  | 0 | 0.96883 | 0.03117 | group_6852 | Isocitrate lyase |
|  | 3277 | 1 | 0.35969 | 0.64031 |  | 0 | 0.96819 | 0.03181 | *thiG* | Thiazole synthase |
|  | 3284 | 1 | 0.35969 | 0.64031 |  | 0 | 0.96819 | 0.03181 | group_5996 | Imidazole glycerol phosphate synthase subunit HisH |
|  | 3586 | 1 | 0.13975 | 0.86025 |  | 0 | 0.95545 | 0.04455 | group_5655 | Isocitrate dehydrogenase [NADP] |

1 – Maximum likelihood posterior probability that this COG was absent at this site in the inferred ancestral reconstruction. Genes were called as absent if p\_0 >= 0.95.
2 - Maximum likelihood posterior probability that this COG was present at this site in the inferred ancestral reconstruction. Genes were called as present if p\_0 >= 0.95.
3 – Predicted names and functions of COGs of interest were recovered from the associated Roary matrix. Sequential COGs are not necessarily syntenic in any given genome.

### *Kingella* species are unified by gene presence/absence in the pangenome

Finally, another recent evolutionary study of these taxa has used less stringent similarity and gene presence cutoffs to define the core genome for species in the *Neisseriaceae* for subsequent phylogenetic analyses (34). We replicated this analysis by reanalyzing the data using a sequence similarity cut off of 50% for genes present in 80% of analyzed genomes. By lowering the threshold for a gene to be considered part of the core genome, we increase the number of genes considered from 109 at the 75% similarity level up to 620 genes. Notably, as we increased the number of genes considered part of the core genome by decreasing the stringency of our Roary analysis, the presence of a polyphyletic *Kingella* clade remains constant (Fig. 4A), demonstrating the robust relationships between these genera that are obtained when we consider the core genome of species in the *Neisseriaceae*.

**Figure 4.**
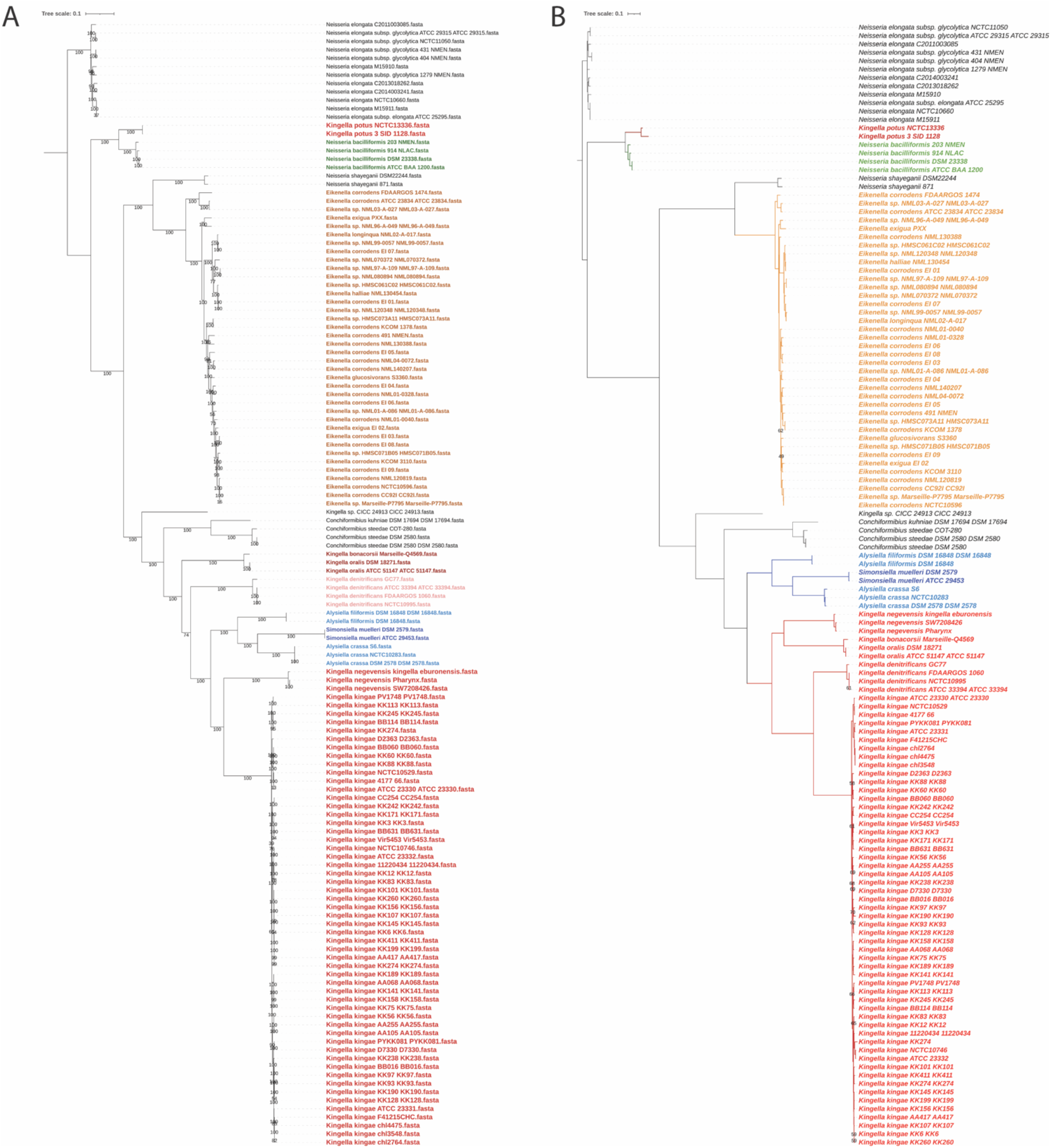
**A**. The core genome of the strains from the 13 species most closely related to *Kingella kingae* was determined using Roary with a minimum homology cutoff of 50% with COGs present in at least 80% of the species. The core genomes, which encompass approximately two million sites, were aligned with MAFFT and used to reconstruct a maximum likelihood phylogeny. As above, *K. potus* is only distantly related to the rest of the species in the genus *Kingella*. Additionally, the genera *Alysiella* and *Simonsiella* fall within the *Kingella* clade, suggesting the core genome of the species in each of these genera is very closely related to that of *Kingella*. Bootstraps greater than 70% are shown. The phylogeny is rooted by the *Neisseria elongata* clade. **B**. A binary gene presence/absence matrix was generated using Roary at the 75% similarity cut-off, and includes both the core and accessory genes of all species. A phylogenetic tree based on this matrix was constructed using RAxML via the GAMMA model of rate heterogeneity. This analysis, based solely on gene content unites *Kingella sensu stricto* into a monophyletic group. Bootstraps greater than 70%are shown. The phylogeny is rooted by the *Neisseria elongata* clade.

To better understand how the pangenome impacts the relationships described in the above analyses, we reconstructed the phylogeny considering only gene presence and absence in the pangenome. This analysis was performed with a binary COG presence/absence matrix as defined by Roary with a 75% similarity cut-off with phylogenetic inference in IQ-Tree. For the first time in any of our analyses, all named *Kingella* species except for *K. potus* are monophyletic, excluding the *Simonsiella* and *Alysiella* genomes. To identify genes that drive the unification of *Kingella sensu stricto (Kingella s.s.*), IQ-Tree was used to perform ancestral reconstructions of the species in this tree. These ancestral reconstructions facilitate the identification of the genes that drive the unification of *Kingella* into a monophyletic clade. Discounting statistically ambiguous sites, monophyly is driven by the loss of eight genes in *Kingella s.s*. (Table 4), all of which fall in the accessory genome. There are no gene gains that support this clade, and therefore this monophyly is not well supported.

These data further suggest that horizonal gene transfer (HGT) can confound our ability to define species relationships and that limits on HGT may be important to consider when defining species in the *Neisseriaceae*. HGT is driven by DNA uptake sequences (DUS) in the *Neisseriaceae* (42–44), which are largely thought to be species specific. The DUS in the CASK clade have been previously characterized (42). Type genomes for each species were used to define DUS preference by word search. *Alysiella filliformis*, *Alysiella crassa*, *Simonsiella muelleri*, and *Kingella oralis* genomes are enriched for the AG-SimDUS (AGGCTGCCTGAA) and the AG-KingDUS (AGGCAGCCTGAA). *Kingella kingae, Kingella negevensis*, and *Kingella denitrificans* genomes are enriched for the AA-King3DUS (AAGCAGCCTGCA). *Conchiformibius steedae* and *Conchiformibius kuhniae* do not appear to use any of these previously defined DUS.

### The CASK species are unique based on cellular morphology and lifestyle

To identify the genes that differentiate the species in the CASK clade and to define clusters of orthologous groups (COGs) unique to groups of interest, Scoary was used to analyze the pangenome analysis generated by Roary at the 75% sequence similarity level. The presence of genes is defined as diagnostic for the in-group if they are 100% specific, are 100% sensitive, and have a *p* > 0.001. These cutoffs reduce the impact of shared mobile genetic elements (MGEs) such as prophages or genomic islands.

First, the CASK clade was compared against the rest of the species included in the phylogenetic analyses (Fig. 5A, table S5). This clade is enriched for 1360 COGs that are absent in the rest of the analyzed species, providing strong evidence supporting the evolutionary relatedness of this group. The average genome size for the species in this group is approximately 2,000 coding sequences (CDS). To understand putative roles of these COGs in the CASK species, COG groups and Gene Ontology (GO) functional annotations were defined with eggnog mapper (Fig. 5B). Approximately half of the COGs were assigned to the “function unknown” group (n = 656). Of the remaining COGs, the majority play putative roles in cell wall, membrane, and envelope biogenesis (n = 276); DNA replication, recombination, and repair (n = 224); or amino acid transport and metabolism (n = 205).

**Figure 5.**
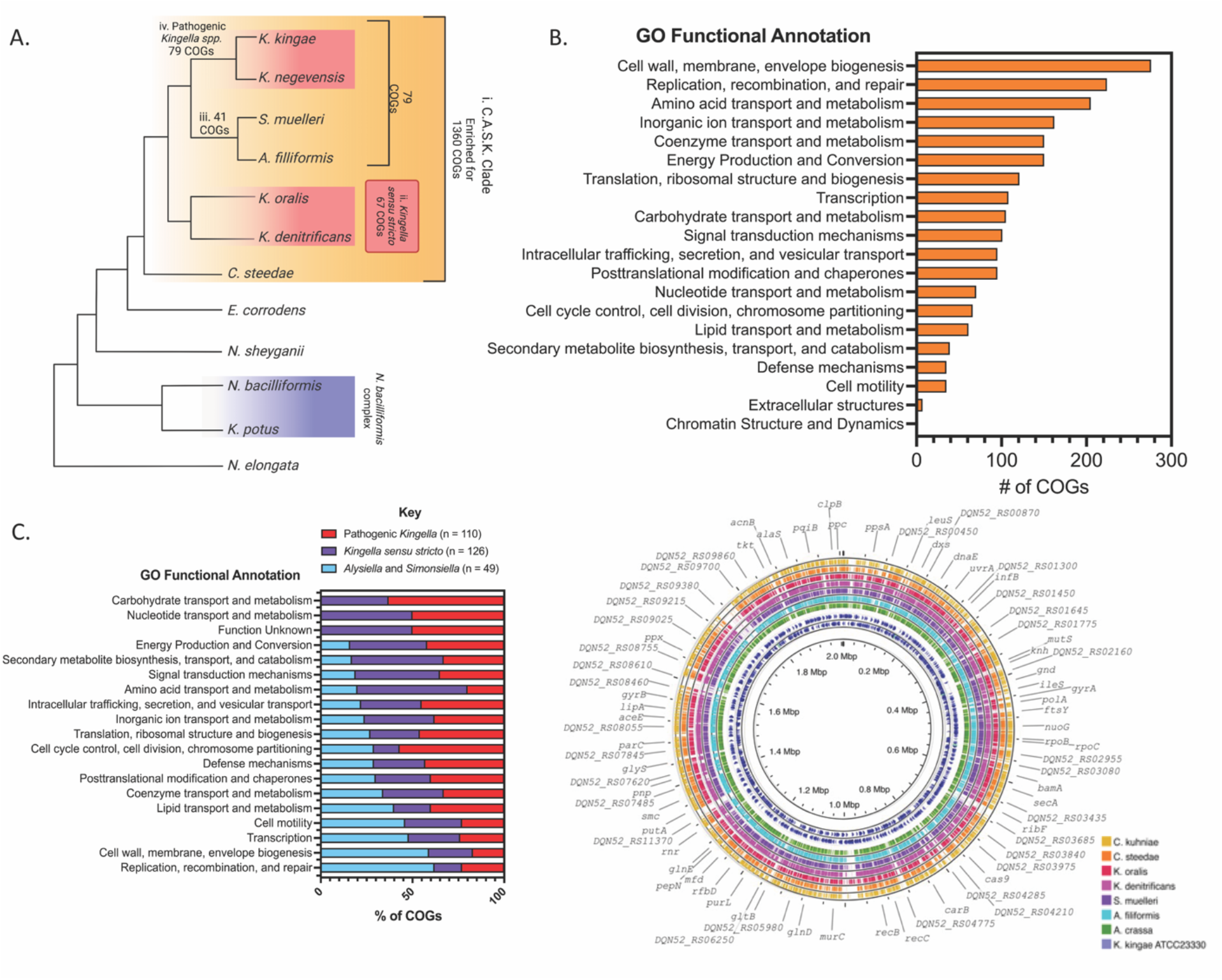
**A**. Scoary was used to generate lists of genes differentially encoded genes comparing (i) CASK species against the *Neisseriaceae*, (ii) *Kingella sensu stricto* (iii) *Alysiella* and *Simonsiella* or (iii) pathogenic *Kingella* against the CASK clade. Finally, *Alysiella* and *Simonsiella* COGs were compared against the pathogenic *Kingella*. Diagnostic COGs are defined as COGs with 100% sensitivity and specificity to the in-group, and filtered to *p < 0.001*. Please see Supplemental tables 5-9 list for full COGs sets. **B-C**. Diagnostic gene sets for each group were annotated using eggNOG mapper, and classified by Gene Ontology functional groups using the NCBI COG database. B. The CASK clade is enriched for unique COGs with putative roles in cell wall, membrane and envelope biogenesis, as well as DNA replication, recombination and repair. COGs with no functional annotations were omitted from this graphic (n = 656 COGs). **C**. Similarly, the majority of the unique COGs present in *Alysiella* and *Simonsiella* may function as part of DNA replication, recombination and repair pathways and cell wall, membrane and envelope biogenesis. *Kingella sensu stricto* and the pathogenic *Kingella* species encode more unique COGs implicated in carbohydrate, amino acid, or nucleotide transport and metabolism. **D**. Type strains of each species in the CASK clade was compared to the *K. kingae* ATCC 23330 genome (inner circle) using BLASTn. Bars in each ring represent regions of sequence similarity between genomes. Each of the CASK species shows significant sequence similarity to nearly all of the *K. kingae* reference, further demonstrating how closely related each of these species are. Regions of dissimilarly map to regions that encode putative MGEs, particularly prophages.

Next, *Kingella sensu stricto (Kingella s.s.*), comprised of *K. oralis, K. denitrificans, K. negevensis*, and *K. kingae*, was compared to the rest of the CASK clade. Only 67 COGs meet the criteria to be diagnostic for *Kingella s.s* (Table S6). When *Alysiella* and *Simonsiella* or the pathogenic *Kingella* spp. were compared to the rest of CASK clade, we identified only 41 or 79 diagnostic COGs, respectively (Fig. 5A). Again, we assigned putative functional roles for the COGs in each of these groups (Fig. 5C). The COGs diagnostic for *Kingella s.s*. and the pathogenic *Kingella* spp (*K. negevensis* and *K. kingae*) are primarily involved in carbohydrate transport and metabolism or nucleotide transport and metabolism. By contrast, *Alysiella* and *Simonsiella* diagnostic COGs are involved in DNA replication, recombination, and repair or cell wall, membrane, and envelope biogenesis (Fig. 5C).

Finally, we compared the genomes of the type-strain for each species in the CASK clade using BLASTn against *K. kingae* ATCC 23330 (Fig. 5D). We found high levels of nucleotide sequence similarity across the *K. kingae* genome relative to the related species. There are few notable regions of dissimilarity, which appear to be MGEs in the *K. kingae* genome.

## Discussion

Classification of the species in the *Neisseriaceae* has been difficult using traditional metrics of relatedness, both phenotypically and phylogenetically. As a result, the family has been restructured several times since its inception to include or exclude genera (15, 24, 36, 45–47). Species in this family resist robust phylogenetic classification due to pervasive horizontal gene transfer and recombination, including in the 16S rRNA sequence (18–20, 42). Due to the large diversity of species and variability, phenotypic classification using strict phenotypic classification is either difficult or impossible without additional sampling. For example, presence of catalase is conserved among *Neisseria* isolates, and yet isolates of *N. elongata* have been recovered that lack the catalase enzyme(33). Furthermore, as more species are proposed using culture-independent methods or based on a single isolate, it will be essential to establish phylogenomic species definitions for the *Neisseriaceae* (1, 18, 20).

In this study, we employed a phylogenetic approach to define the evolutionary relationships within a major subgroup of the *Neisseriaceae*, defined by the genera *Eikenella, Conchiformibius, Alysiella, Simonsiella, Kingella*, and select *Neisseria* species. Examination by 16S rRNA sequence unveiled a very recent common ancestor between *K. potus* and *N. bacilliformis*, which was further supported by several other genome relatedness indices (Fig. 1, Tables 1,3). *K. potus* is a catalase-negative species collected from an infected wound that resulted from a bite by a captive South American Kinkajou (*Potus flavus*) (25). *N. bacilliformis* is an opportunistic pathogen that has been recovered from human infections, varies in the production of catalase (48), and is phenotypically identical to *N. elongata* subsp. *elongata*. These species were reported nearly simultaneously, preventing their direct comparison. Our reanalysis suggests that *K. potus* is much more closely related to *N. bacilliformis* than to any other *Kingella* species, though it is likely a distinct species from *N. bacilliformis*. Inclusion in the genus *Kingella* was likely due to a negative catalase test, underscoring the importance of a genomic approach to classification of these isolates.

The phylogenomic analyses also uncovered close evolutionary relationships between *<u>C</u>onchiformibius* sp., *<u>A</u>lysiella* sp., *<u>S</u>imonsiella* sp., and *<u>K</u>ingella* sp., which we have named the “CASK” clade (Fig. 3). The CASK clade is a major branch of the *Neisseriaceae* that includes both invasive pathogens (*K. negevensis* and *K. kingae*) and oral-associated commensals. *Alysiella* and *Simonsiella* are genera associated with the oral microbiome of non-human mammals (15, 47, 49). *Conchiformibius* spp., *Alysiella* spp., and *Simonsiella* spp. are remarkable for their multicellularity, which is the result of incomplete septation by the forming peptidoglycan during bacterial division (34).

The reproducibility of the core genome phylogenetic structure very strongly supports the close relationship between the species of the CASK clade and the exclusion of *K. potus* (Fig. 3–4). We hypothesized that prior misclassification could be due to high levels of horizontal gene transfer of accessory genes between *Kingella (sensu stricto*) genomes. To test this hypothesis, we generated a presence/absence matrix generated by Roary at 75% sequence similarity level, a technique that emphasizes accessory genes over core genes, and we reconstructed a phylogeny using the RAxML binary model with gamma model of rate heterogeneity and bootstrapped 100 times (Fig. 3B). This was the only analysis in which we observed a monophyletic *Kingella* genus, though *Kingella potus* was still excluded. We favor the analyses that are based on nucleotide sequence of the core genome because they contain much more genomic information (~76,500 variable sites compared to ~29,000 variable genes) and likely lessen confounding signal from recombined areas. A separate whole gene-based analysis also identified more diagnostic genes uniting *Alysiella, Simonsiella*, and the pathogenic *Kingella* species (n=79) than genes uniting the *Kingella* genus *sensu strictu* (n=69). Additionally, as *Simonsiella, K. oralis*, and *K. kingae* utilize specific DNA-uptake sequences (DUS), suggesting they may be able to share DNA more easily among themselves as compared to the rest of the *Neisseriaceae*. Given that members of the *Neisseriaceae* are highly recombinogenic and undergo significant HGT of accessory genes, we believe that the core genome is a higher fidelity evolutionary record for this family (20).

The CASK clade has a large set of diagnostic genes that differentiate these species from the remainder of the *Neisseriaceae*. Interestingly, this group is enriched for processes that impact cell wall and envelope biogenesis (Fig. 5B), likely driven by the unusual cellular morphologies of *Conchiformibius* spp., *Alysiella* spp., and *Simonsiella* spp. These genera are distinctive for being multicellular, longitudinally-dividing (MuLDi) species that comprise the only known animal multicellular bacterial symbionts (34). Nyongesa and co-workers hypothesize that the MuLDi phenotype evolved twice in the *Neisseriaceae*, first in *Conchiformibius* species and then in the common ancestor of *Alysiella* and *Simonsiella*, a hypothesis that is strongly supported by our phylogenetic analyses. This hypothesis is further supported by consideration of the diagnostic genes that differentiate *Alysiella* and *Simonsiella* from the rest of this group (Fig. 5C). We again find an enrichment for COGs implicated in cellular morphology, as we would expect if these genera evolved the MuLDi phenotype independently of *Conchiformibius* spp. By contrast, *Kingella s.s*. is enriched for COGs associated with nutrient transport and metabolism (Fig. 5C).

Our study suggests that the current taxonomic classifications of species in the *Kingella* genus may not reflect evolutionary history. Based on our results, *K. potus* is more related to *N. bacilliformis* than to other *Kingella* species by 16S rRNA, OGRI, and whole genome analyses and should probably be assigned to the genus *Neisseria* or another genus entirely. Additionally, phylogenomic analysis of the core genomes of *Alysiella* and *Simonsiella* indicates their close relationship to the *Kingella* genus and supports their reassignment to *Kingella*, resulting in a monophyletic genus.

## Methods

### Phylogenetic Analysis

840 of the available 16S rRNA gene sequences for isolates from the *Neisseriaceae* and in the RefNR database were downloaded from the SILVA r138.1 database on September 08, 2022 (50, 51). 16S sequence accession numbers are listed in table S1. 16S rRNA sequences were aligned with MAFFT, and a phylogeny was reconstructed using RAxML v. 8.4.2 with the general time reversible model of nucleotide substitution and gamma model of rate heterogeneity, and bootstrapped 100 times (52, 53).

Whole genome sequences for *Neisseria elongata, Neisseria sheyganii, N. bacilliformis, Eikenella corrodens, Alysiella filiformis, Alysiella crassa, Simonsiella muelleri, Conchiformibius steedae, K. oralis, K. bonacorsii, K. potus, K. denitrificans, K. kingae*, and *K. negevensis* were downloaded from the NCBI Genome database on September 09, 2022. Sequence accession numbers are listed in table S2. In total, 134 genomes were reassembled with Prokka v1.14.6, using default settings for bacterial sequences (54). Phenotypic characteristics for each of these species were collected by literature review and are listed in table S3. Assemblies were analyzed with Roary v3.13.0. to identify the core genome (40). Core genes were clustered at the 75%, 80%, 85%, 90%, and 95% sequence similarity level and were limited to only those clusters of orthologous groups (COGs) found in all genomes (-cd 100), unless otherwise noted. Additionally, other recent evolutionary study of these taxa has used less stringent similarity and gene presence cutoff to define the core genome for species in the *Neisseriaceae* for subsequent phylogenetic analyses (8, 34). We replicated this analysis by reanalyzing the data using a sequence similarity cut off of 50% for genes present in 99% or 80% of analyzed genomes. Nucleotide sequences of core genes were compared in a codon-aware alignment by PRANK within Roary, and core gene alignments were used to reconstruct phylogenies using RAxML as above. Phylogenetic trees were annotated using Figtree v.1.4.4 and ITOL (55). Bootstrap values >70 are shown.

To calculate phylogenetic tree similarity, final trees were analyzed in R using the Ape v5.6-2 (56), ggtree v3.1.2. (57), and TreeDist v.2.5.0 (58, 59) packages. The Robinson-Fould distance and normalized Nye Simillarity Score were determined for each pair of trees (58–61).

Ancestral reconstructions were performed using the Roary gene presence/absence (PA) matrix generated at the 75% sequence similarity cut-off. The matrix was converted into a FASTA formatted file of binary data using R. This file was used for phylogenetic reconstruction in IQ-Tree v 1.6.12 with automatic model calling (62, 63). Branch support was determined via ultrafast bootstrapping (-bb 1000) (64), and ancestral states (-asr) were calculated with default cut-offs. Full inferred ancestral state matrices are available in Supplementary File 2. To determine the COGs that unite *Kingella s.s*., the inferred PA matrix of common ancestor of *Alysiella* spp., *Simonsiella* spp., and *Kingella s.s*. (Node 7), was compared to the inferred PA matrix of common ancestor of *Kingella* spp. Statistically ambiguous sites present in either PA matrix were removed from this analysis.

### Gene Set Enrichment

Scoary v 1.6.16 was used to further analyze the gene presence absence matrix generated by Roary at the 75% sequence similarity cutoff (65). The -r flag was used to limit the set of strains considered in this analysis. COGs were eliminated from this analysis if the Bonferroni-corrected p-value >0.0001. Diagnostic genes are only those genes which 1) meet this statistical threshold, 2) are enriched in the clade of interest, and 3) have specificity and sensitivity of >80%.

To identify pathways that are enriched in each phylogenetic group, the representative COG sequence was collected from the Roary output files and compiled into a gene set. Gene sets were analyzed with the COG database using eggNOG-mapper v 2.1.9 (66, 67), with a DIAMOND alignment (68) and default settings for bacterial species (Table S4-9). Results were plotted using Graphpad Prism.

### Genome Relatedness Indices

Average nucleotide identity (ANI) and digital DNA-DNA hybridization (dDDH) between the type strains of these species using FastANI v.1.33 and the Type Strain Genome Server (TYGS), respectively (69–71).

## Abbreviations

ANI: Average Nucleotide identity
dDDH: digital DNA-DNA hybridization
OGRI: overall genome relatedness index
COGs: clusters of orthologous gene groups
MuLDi: Multicellular, longitudinally-dividing
DUS: DNA-uptake sequences
MALDI-TOF MS: Matrix-assisted laser desorption-ionization time of flight mass spectrometry
CA: common ancestor

